# cProSite: A web based interactive platform for online proteomics, phosphoproteomics, and genomics data analysis

**DOI:** 10.1101/2023.06.10.543932

**Authors:** Dunrui Wang, Xiaolan Qian, Yi-Chieh Nancy Du, Beatriz Sanchez-Solana, Kailing Chen, Madhu Kanigicherla, Lisa M. Jenkins, Jason Luo, Sarah Eng, Brian Park, Ben Chen, Xiaozheng Yao, Thuong Nguyen, Brajendra K. Tripathi, Marian E. Durkin, Douglas R. Lowy

**Author notes:** Correspondence: Douglas R. Lowy or Dunrui Wang NCI, NIH, Building 37, Room 4106, Bethesda, MD 20892.

## Abstract

We developed cProSite, a website that provides online genomics, proteomics, and phosphoproteomics analysis for the data of The National Cancer Institute’s Clinical Proteomic Tumor Analysis Consortium (CPTAC). This tool focuses on comparisons and correlations between different proteins and mRNAs of tumors and normal tissues. Our website is designed with biologists and clinicians in mind, with a user-friendly environment and fast search engine. The search results of cProSite can be used for clinical data validation and provide useful strategic information to identify drug targets at proteomic, phosphoproteomic, or genomic levels. The site is available at http://cprosite.ccr.cancer.gov.

**Significance:** An interactive database for the expression and correlation for proteins, proteomic phosphorylations, and mRNA levels has been developed for analyzing the molecular alterations between tumors and normal tissues of various tumor types.

## Introduction

The National Cancer Institute’s Clinical Proteomic Tumor Analysis Consortium (CPTAC) is part of a national effort to accelerate the understanding of the molecular basis of cancer through the application of large-scale proteomic and genomic analysis, or proteogenomics (1). We developed a web-based interactive platform, **C**ancer **Pro**teogenomic Data Analysis **Site** (cProSite), that provides on-line proteomics, phosphoproteomics, and genomics analysis for the CPTAC data. Compared with regular analytical methods, this platform analysis has several advantages, including: 1) faster analysis online, 2) more user-friendly environment, and 3) less need for bioinformatics expertise to perform the analysis. In this study, we show examples of analysis from cProSite to demonstrate its utility.

## Materials and Methods

### Data requirement and web platform development

The current cProSite includes 12 tumor types (brain cancer, breast cancer, colon cancer, head and neck cancer, kidney cancer, liver cancer, lung adenocarcinoma, lung squamous carcinoma, ovarian cancer, pancreas cancer, stomach cancer, and uterine cancer).

The proteomics and phosphoproteomics data from cProSite are derived from the Proteomic Data Commons (PDC, https://proteomic.datacommons.cancer.gov/pdc/) and CPTAC portal (https://proteomics.cancer.gov/data-portal). The raw data of CPTAC from various tumor types have been processed uniformly through the CPTAC Common Data Analysis Pipeline (CDAP), enabling comparisons between different samples for individual cancer types (2). The proteomics data files with the names “Proteome.tmt11.tsv” or “Proteome.tmt10.tsv” are selected. The phosphorylation data files with the name “Phosphoproteome.phosphosite.tmt11.tsv or Phosphoproteome.phosphosite.tmt10.tsv” are selected. The relative abundances of the proteins and phosphorylations in CPTAC are presented as TMT log2 ratios computed by the CDAP. For proteomics data, shared values are chosen. The stomach data were an exception, as they used values of tumor/normal pairs, and the web platform adjusted the presentation accordingly. CPTAC mRNA data from cProSite are derived from the Genomic Data Commons Data Portal (https://portal.gdc.cancer.gov/). TCGA data manifest file are from https://docs.gdc.cancer.gov/Data/Release_Notes/Data_Release_Notes/#data-release-330 version 31 and HTseq datasets are download using the GDC DTT. A new dataset of cProSite will be updated upon the new release of CPTAC.

Data derived from assembly files were generated to load into the database SQLite table (SQLite 3.40.0). React, a JavaScript library for building web interfaces, was used to create cProSite’s front-end site, with a NodeJS backend to query the database. Because of mass spectrometry and analysis algorism limitations, not all proteins and real phosphorylation sites are detected in individual samples. The codes of web platform used in this manuscript are available at https://github.com/CBIIT/nci-webtools-ccr-cprosite.

## Results

### cProsite to study differences between tumors and normal tissues in protein abundance, mRNA level, phosphorylation level and phosphorylation level per protein abundance

cProSite allows users to perform analysis for protein abundance and phosphorylation levels between tumors and normal tissues (normal tissues adjacent to the tumor (NAT) in most tumor types). Protein abundance is comparable in individual tumor type (TMT log2 ratio) and among various tumor types (log2 fold changes between tumors vs normals). Figure 1A shows one example, log2 fold changes in caveolin 1 (CAV1) protein between tumors and normal tissues among the 12 tumor types. The result shows that CAV1 is downregulated in all the tumor types selected except for kidney cancer. cProSite provides a ‘tumor view’ to view the differences in protein abundance between tumors and controls in unpaired samples (box blot) or paired samples (bar plot, CAV1 downregulated in lung adenocarcinoma, Figure 1B).

**Figure 1.**
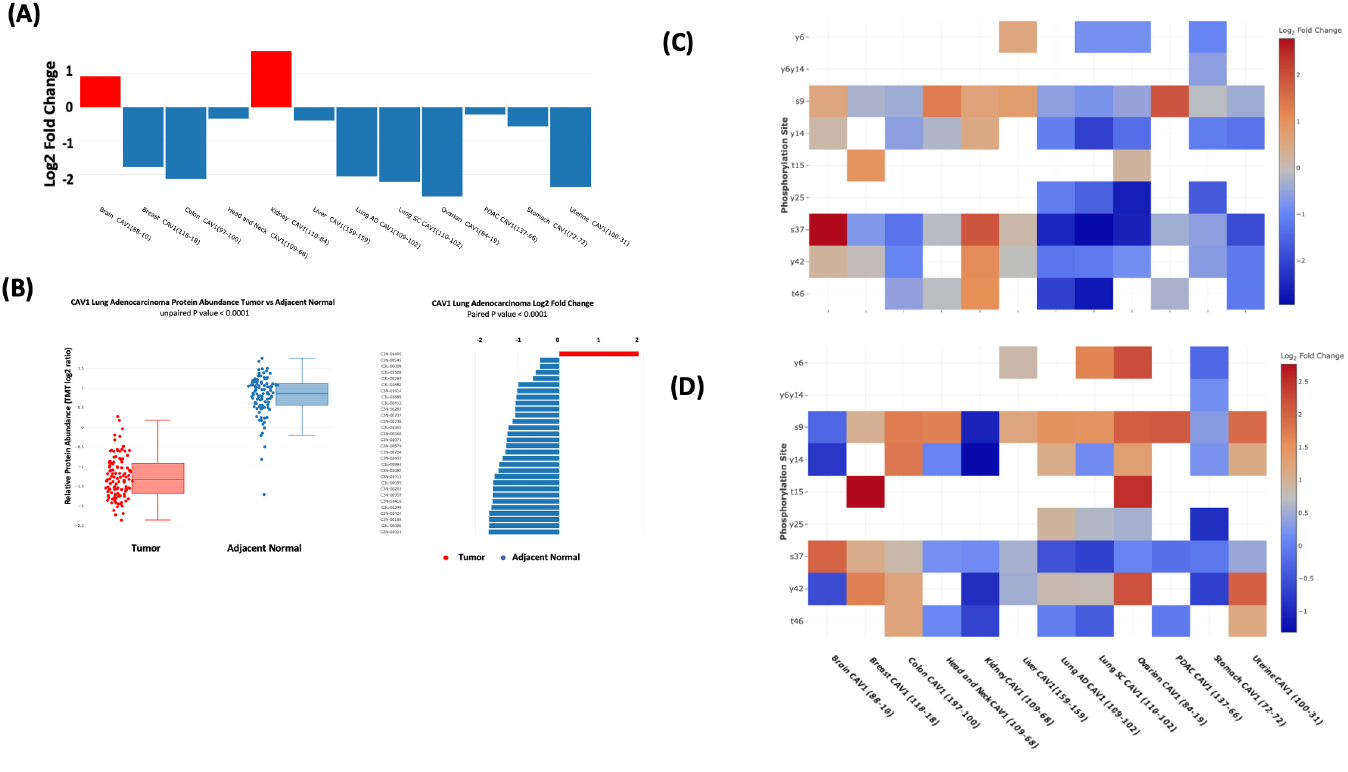
Comparison between tumors and normal tissues for protein abundance and phosphorylation level by cProSite. (A) log2 fold change of CAV1 protein abundance between tumors and normal tissues from 12 tumor types. (B) Box p ot comparison (unpaired) and bar plot of log2 fold change of CAV1 between tumor 2nd paired NAT in lung adeneeareinema for protein abundance. (C) Heatmap shewing die log2 fold changes of multiple CAV1 phosphorylation sites in various tumor types. (D) Hcatmap showing the log2 fold changes of normalized phosphorylation level on the same set of sites of CAV1 (phosphorylation site level / protein abundance) in various tumor types.

For comparison of phosphorylation levels from individual phosphorylation sites in various tumor types, the cProSite ‘summary page’ for a protein phosphorylation has a heatmap-like presentation with various tumor types. Figure 1C shows levels of phosphorylation sites of CAV1, in which the log2 fold changes of phosphorylation sites in lung cancers are negative. The changes are altered for some phosphorylation sites if the concept of phosphorylation level per protein abundance level is introduced (Figure 1D). The results represent the actual phosphorylation level of CAV1 per protein level since overall phosphorylation level in a specific site is not necessarily proportional to the protein level.

cProSite also allows users to perform analysis for mRNA levels between tumors and normal tissuess, and users can compare expression of the same gene at both protein and RNA levels.

### cProSite to analyze the correlation between protein abundance and RNA expression of the same gene and between protein/phosphorylation levels of different genes

Exploring the relationship between protein and mRNA expression in tumors and normal tissues is a featured function of CPTAC datasets. The relationship between protein and mRNA levels in the cells under various scenarios - such as steady state, long-term state changes, and short-term adaptation - demonstrates the complexity of gene expression regulation, especially during dynamic transitions (3). The patterns of different correlations for the genes may be completely different. Some important proteins involved in cell cycle G2/M transitions are expressed at relatively low mRNA levels in normal tissues with poor correlation to protein levels (Figure 2A-B). In tumors, both mRNA and protein levels are greatly enhanced and have an increased correlation, suggesting that active protein synthesis is coupled well with increased mRNA transcription in tumors.

**Figure 2.**
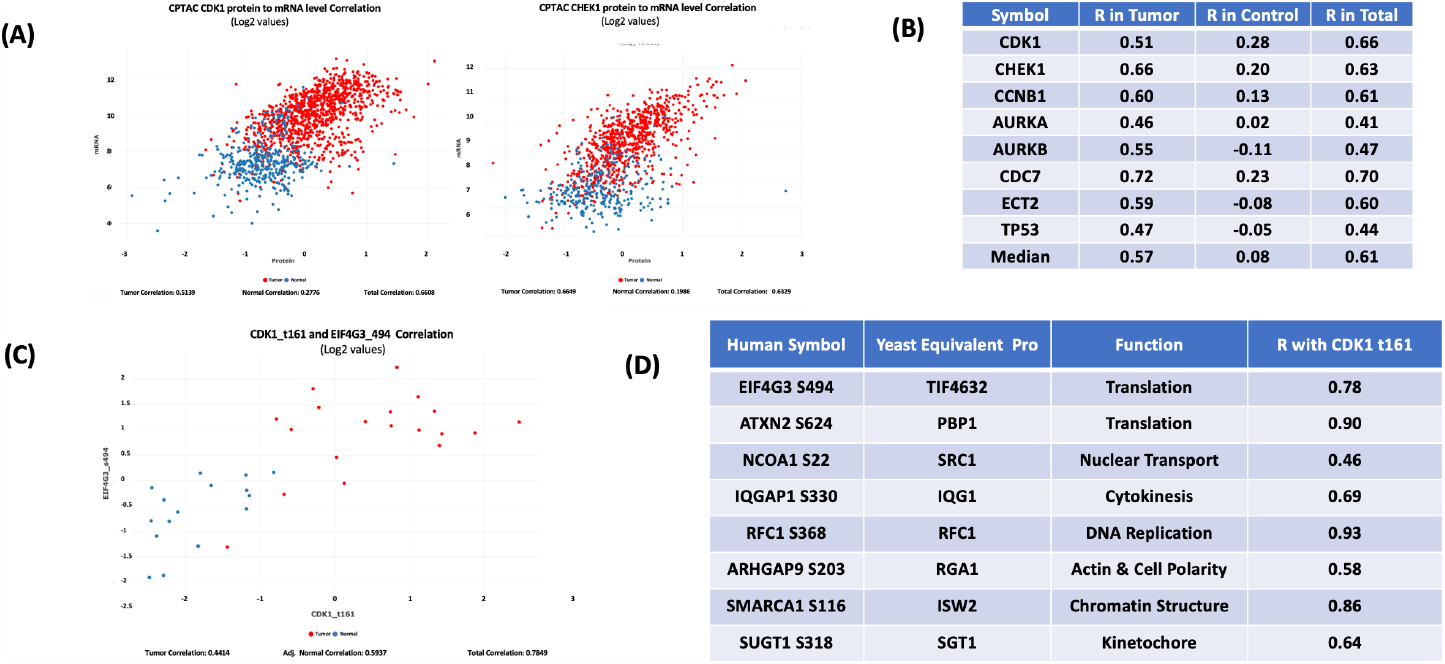
Correlation among protein abundance, phosphory lation site level, and mRNA level by eProSitc. (A) The correlation of protein and mRNA levels of important cell cycle related protein CDK1 and CHEK1 from combined 11 tumor type datasets (brain, breast cancer, colon cancer, head and neck cancer, kidney cancer, liver cancer, lung adenocarcinoma, lung squamous carcinoma, ovarian cancer, pancreas cancer, and uterine cancer). (B) List of cell cycle proteins with high correlation between protein and mRNA from combination of 11 tumor types. (C) The correlation between CDK1 t161 and TIF4G3 S494 expression in lung adenocarcinoma. (D) List of the correlation between CDK1 t161 and phosphorylation sites of CDK1 substrates predicted from yeast homologues in lung adenocarcinoma.

cProSite provides search tools that evaluate the correlation between protein abundance in the following manner: 1) protein versus protein; 2) protein versus phosphorylation site; 3) phosphorylation site versus phosphorylation site. Phosphorylation site abundance can also be presented as phosphorylation site per protein level; 4) mRNA versus mRNA.

CDK1 is a serine/threonine kinase that drives the G2/M phase of the cell cycle, and increased CDK1 kinase activity is a frequent feature of all tumor types. More than 500 phosphorylation sites from more than 300 CDK1 substrates have been identified in yeast (4). Figure 2C-D shows the correlation between the CDK1 active site threonine 161 level and the predicted phosphorylation site level of the yeast homolog, including EIF4G3 (translation), ATXN2 (translation), NCOA1 (nuclear transport), IQGAP1 (cytokinesis), RFC1 (DNA replication), ARHGAP9 (actin & cell polarity), SMARCA1 (chromatin structure), and SUGT1 (kinetochore) in human lung adenocarcinoma. The results suggest that phosphorylation levels of those proteins associated with CDK1 t161 are well correlated in both normal and tumor stages, with enhanced CDK1 activity and high phosphorylation of its substrates. These correlations may be useful for identifying prognostic markers.

### cProSite to validate clinical data and confirm biological experiment results

The protein abundance changes between tumor and NAT from CPTAC are good indicators that can be used for clinical data validation. We have previously reported that expression of the receptor for hyaluronic acid mediated motility (RHAMM, gene name: HMMR) is associated with poor prognosis and metastasis in non-small cell lung carcinoma (NSCLC) (5). Tissue microarray (TMA) analysis that evaluated the expression of RHAMM in NSCLC revealed positive staining for more than one-half of primary and metastatic tumors (Figure 3A). The results are consistent with the cProSite protein abundance comparison between tumors and NATs in lung adenocarcinoma and lung squamous cell carcinoma, in which RHAMM/HMMR is highly expressed in tumors (Figure 3B-C).

**Figure 3.**
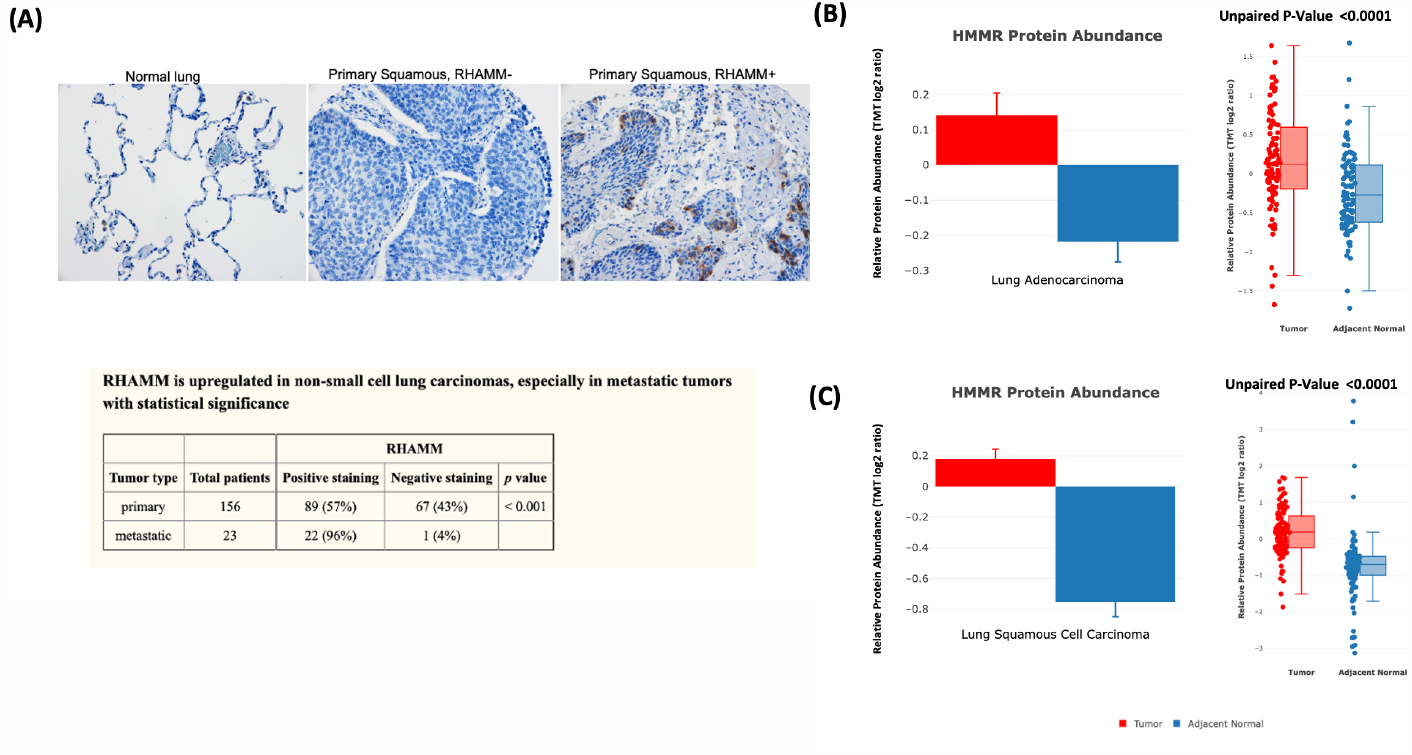
Using cProSite to confirm RHAMM/HMMR upregulation in non-small cell lung carcinoma tissue microarray. (A) Tissue microarray detection of RHAMM/HMMR in NSCLC. The results are derived from Figure 1 and Table 3 of our published work (5). (B) Comparison of RHAMM/HMMR protein abundance between tumor and NAT in lung adenocarcinoma. (C) Comparison of RHAMM/HMMR protein abundance between tumors and NAT in lung squamous cell carcinomas.

To study a possible direct effect of CDK1-dependent phosphorylation on a protein involved in cell cycle progression, we selected Epithelial Cell Transforming 2 (ECT2), which is a Rho family-specific GEF (guanine nucleotide exchange factor) with a role in G2/M (6). In agreement with the proteomic data analysis, the mass spectrometry analysis revealed increased phosphorylation at multiple sites of ECT2, including T359, T373, T444, S861, and S889 in 293T cells synchronized at the G2/M phase. The phosphorylation was observed to decrease upon treatment with the CDK1 inhibitor RO-3306 (Figure 4). Biological experiments confirmed that phosphorylation of these sites enhanced the pro-oncogenic and GEF activity of ECT2 (7).

**Figure 4.**
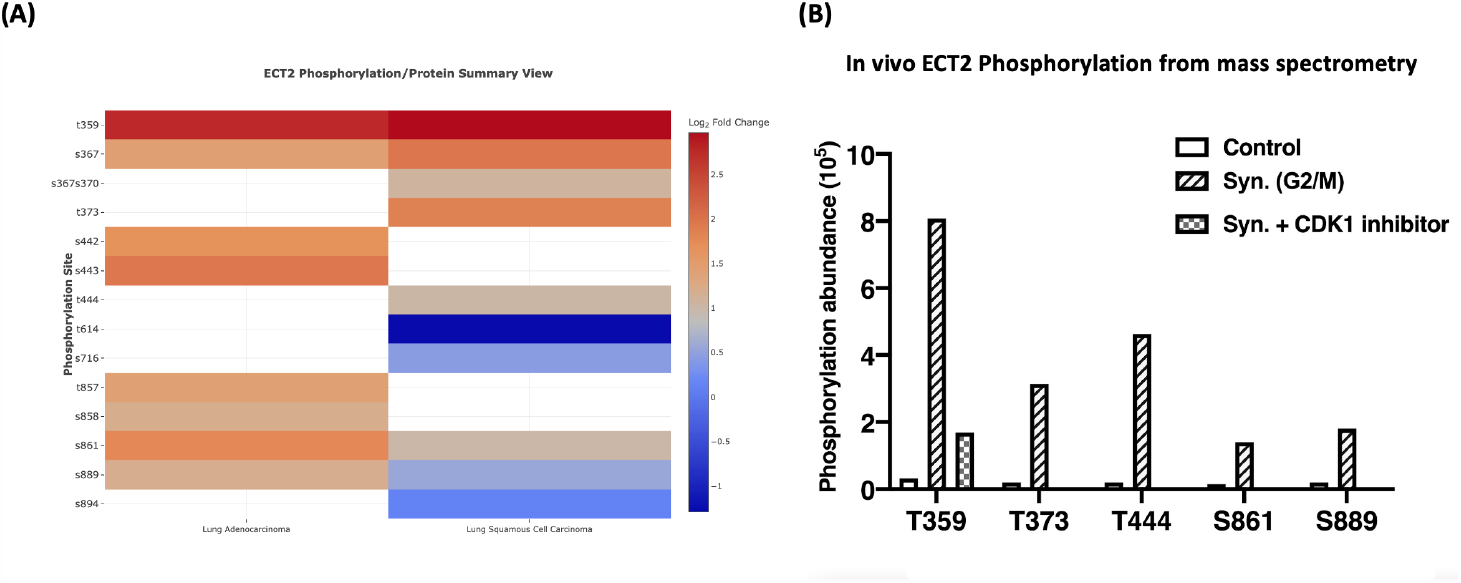
Characterization of phosphorylation sites of ECT2 using CPTAC datasets. (A) ECT2 phosphorylation site log2 fold changes for lung adenocarcinoma and lung squamous cell carcinoma. (B) Validation of phosphorylation sites of ECT2 T359, T373, T444, S861, and S889 using human kidney epithelial cell line 293T. The cells were synchronized by Nocodazole to reach the G2/M phase and treated with or without CDK1 inhibitors afterwards. The selected phosphorylated residues in purified ECT2 from these cell extracts were detected by mass spectrometry (MS) analysts and plotted by GraphPad Prism software.

## Discussion

Various computational packages for CPTAC make data and tools available and accessible to the greater research communities such as UALCAN (8), cBioPortal (9), and LinkedOmics (10). UALCAN (8) uses Z scores to compare CPTAC’s protein abundance and phosphorylation levels from various tumor types in a web platform-based Pan-cancer global proteomic data analysis. cBioPortal (9) provides a function to display the co-expression of two different proteins, or protein abundance & mRNA levels from same gene. LinkedOmics (10) provides a function to display the correlation of the protein abundance between one protein and others that are positively or negatively correlated. The Office of Cancer Clinical Proteomics Research (OCCPR) from the National Cancer Institute has curated a collection of computational tools developed and/or utilized by CPTAC for the processing and analysis of proteogenomic data (11). These tools are useful for analyzing and/or visualizing large-scale proteomics, phosphoproteomics, and genomics datasets from various tumor types.

cProsite has unique features that display 1) the changes between tumor and NAT in various tumor types in levels of protein, protein phosphorylation, and mRNA, 2) Correlation of expression between 2 protein abundance or 2 mRNA levels, 3) Correlation of expression levels between protein and mRNA from the same gene, 4) Correlation among the protein and phosphorylation site levels. Fold changes of protein abundance, phosphorylation levels, and mRNA levels between tumors and normal tissues from genes displays the differences among various tumor types (Figure 1A & 1C). Paired sample comparisons for a specified gene from an individual sample shows the distribution of all the fold changes in the individual tumor type (Figure 1B). cProsite uses phosphorylation levels per protein abundance instead of phosphorylation level alone for the protein phosphoproteomics data analysis, to reflect real phosphorylation up or down regulation in the tumor.

cProSite focuses on the display of the differences in correlation analysis between tumors and normal tissues either in the abundance of two proteins, in mRNA levels of two genes, or in mRNA and protein abundance. In normal tissues, low expression levels and poor correlation between mRNA and protein levels of cell cycle-related genes suggest that tumorigenesis is associated with changes in the regulation of protein abundance (Figure 2A-B).

We used cProSite data analysis to validate our previously published experiment results from non-small cell lung carcinoma TMA (Figure 3) and characterize the protein phosphorylation sites of a particular protein (Figure 4). In summary, cProSite is expected to facilitate investigation of the molecular alterations in various tumor types and the correlation among protein, post translational modification, and mRNA levels.

## Authors’ contribution

**D. Wang:** Conceptualization, resources, formal analysis, validation, investigation, methodology, writing-original draft, project administration, writing-review and editing. **X. Qian, Y.-C. N. Du:** Conceptualization, resources, formal analysis, validation, investigation, methodology, writing-original draft, writing-review and editing. **K. Chen, B. Park, B. Chen, M. Kanigicherla, X. Yao, T. Nguyen:** web platform development. **B. Sanchez-Solana, B. K. Tripathi, M. E. Durkin:** Validation, methodology, writing-review and editing. **L. M. Jenkins:** Proteomics experiment design and processing. **J. Luo, S. Eng:** Data collection, validation, and processing. **D. R. Lowy:** Conceptualization, resources, supervision, funding acquisition, investigation, writing-original draft, writing-review and editing.

## Acknowledgements

This project was supported by office of The National Cancer Institute’s Clinical Proteomic Tumor Analysis Consortium (CPTAC). We thank Dr. Henry Rodriguez, Ana Robles, An Eunkyung, and Xu Zhang from the CPTAC office for critical review and helpful recommendations for this web platform. We thank Drs. Nathan Edwards from Georgetown University, Paul Rudnick from Spectragen Informatics and Mike MacCoss from University of Washington for their helpful discussion regarding CPTAC data analysis.

